# Reduced Pupil-Linked Arousal Marks Implicit Memory for Complex Auditory Sequences

**DOI:** 10.64898/2026.01.27.701958

**Authors:** Wing Shuen Leung, Lawrence Yeung, Mert Huviyetli, Roberta Bianco, Maria Chait

## Abstract

Observers rapidly form long-lasting implicit memories for recurring sensory patterns, supporting adaptive behaviour through learning of environmental contingencies. Here we tested how such memories interact with neuromodulatory systems that regulate arousal—whether implicit familiarity increases arousal through retrieval-related processes or instead decreases arousal via more efficient sensory processing.

We examined these competing accounts using complex auditory tone-pip sequences in which a subset of patterns recurred approximately every three minutes. One group of participants (N=30) actively detected transitions from random to regular structure, whereas a second group (N=40) heard the same sequences while performing a decoy task. Consistent with prior work, recurring regular sequences elicited faster reaction times in the active group, confirming the formation of implicit memory.

Subsequently, both groups listened to previously encountered regular patterns (REGr) and matched novel regular patterns (REGn) while pupil diameter was recorded. Across groups, REGr sequences evoked significantly smaller pupil dilations than REGn sequences, indicating attenuated arousal for familiar patterns. This reduction was sustained until sequence offset in the active group but was transient in the exposure group, suggesting that task demands during exposure shape the stability of pupil-linked memory expression. Importantly, the REGr–REGn pupil difference did not correlate with explicit post-session recognition, demonstrating that the effect was not driven by conscious awareness.

Together, these findings identify pupil dynamics as a robust physiological marker of learned auditory structure and show that implicit auditory sequence memory attenuates arousal. The results support predictive processing accounts in which familiarity reduces uncertainty and, consequently, neuromodulatory drive.

**Significance Statement:** Humans rapidly acquire implicit memories for recurring sounds, yet how such memories influence arousal during subsequent exposure remains poorly understood. Using pupillometry, we show that implicitly remembered auditory sequences evoke reduced pupil-linked arousal relative to matched novel sequences, even when listeners are unaware that the patterns have been previously experienced. This effect dissociates from explicit recognition, which typically yields the opposite “old/new” pattern of larger pupil responses to familiar items. Together, these findings establish pupil dynamics as a sensitive, non-invasive marker of implicit auditory memory and support predictive processing accounts in which familiarity reduces uncertainty and, consequently, neuromodulatory drive.

## Introduction

A fundamental mechanism underlying adaptive behaviour is the brain’s capacity to rapidly form long-lasting implicit memories for recurring sensory patterns, even in the absence of explicit awareness. This process allows the nervous system to extract statistical regularities in the environment, enabling the learning of predictive contingencies that constrain perception and guide action (Robertson, 2007; Schacter et al., 1993).

In the auditory modality, sensitivity to recurring structure has been documented for a variety of stimuli (Bastug et al., 2026; Bianco et al., 2020; Kang et al., 2017; Loui et al., 2010; Saffran et al., 1999). A landmark study by Agus et al. (2010) showed that listeners performing a repetition-detection task formed stable long-term memories for recurring noise snippets, expressed as enhanced detection relative to novel sounds and persisting for up to four weeks (see also Agus & Pressnitzer, 2021; Luo et al., 2013). This learning was attributed to the encoding of idiosyncratic spectrotemporal “noise profiles” and the rapid emergence of neural selectivity in auditory cortex. Extending this work to more constrained stimuli, Bianco et al. (2020) demonstrated robust, incidental learning of recurring tone-pip sequences that differed only in temporal order. This memory generalized across age groups (Bianco et al., 2023) and persisted for up to six months. Because novel and reoccurring patterns differed only in the precise ordering of identical elements, these findings demonstrate that the auditory system can encode and retain fine-grained temporal structure even when patterns are highly similar.

Such robust memory for seemingly meaningless acoustic patterns likely reflects a fundamental operating principle of the auditory system. Rather than serving only explicit behavioural goals, auditory memory may be adaptively tuned to detect sparse repetitions in natural environments, such as recurring animal vocalizations or other biologically relevant sounds, where the rapid recognition of familiar patterns can convey survival-critical information.

How, then, do such implicit memory traces influence the processing of familiar sounds? Remembered noise (Andrillon et al., 2015) and tone patterns (Bianco et al., 2026; Magami et al., 2025) elicit earlier neural responses than novel patterns, even in naïve or distracted listeners, indicating facilitated sensory processing. Source-level analyses further reveal reduced engagement of auditory and frontal networks typically involved in the detection of regularity (Bianco et al., 2026), suggesting a transition from resource-demanding computation to automatic recognition once a sound pattern becomes familiar.

Building on this framework, the present study examines how implicit auditory memory shapes physiological markers of arousal. Pupil diameter provides a time resolved non-invasive index of neuromodulatory activity, reflecting coordinated signaling within the locus coeruleus–norepinephrine and basal forebrain–acetylcholine systems (Aston-Jones & Cohen, 2005; Grujic et al., 2024; Joshi et al., 2016; Joshi & Gold, 2020; Murphy et al., 2014; Reimer et al., 2016). Despite growing interest in pupillometry as a tool for probing cognitive function, the mechanisms linking implicit memory to pupil dynamics remain poorly understood.

Previous work has primarily linked pupil responses to declarative memory, most notably in old/new recognition paradigms. Larger pupil dilations during encoding reliably predict superior subsequent recall, consistent with arousal-dependent memory formation (Kafkas & Montaldi, 2011; Micula et al., 2022; Naber et al., 2012; Pitem & Mama, 2025; Robison et al., 2022). During recall, previously encountered stimuli typically elicit larger pupil dilations than novel ones, a phenomenon known as the pupil old/new effect, often interpreted as reflecting retrieval effort or heightened arousal associated with remembered information (Kafkas, 2024; Lapteva & Martarelli, 2024; Võ et al., 2008), but see (Gomes et al., 2021). While robust (Lapteva & Martarelli, 2024), these effects have been argued to depend on controlled, attention-dependent memory processes (e.g. (Brocher & Graf, 2016, 2017), leaving open the question of whether pupil dynamics can index automatic, incidental memory.

Recent findings suggest that pupil responses are also tightly coupled to implicit adaptive learning and predictive processing (Krishnamurthy et al., 2017; Nassar et al., 2012). In auditory tasks, regular and predictable sound sequences elicit reduced pupil dilation relative to random sequences, consistent with lower arousal under conditions of reduced uncertainty (Huviyetli & Chait, 2026; Milne et al., 2021). From a predictive processing perspective, novel patterns violate expectations and generate prediction errors that drive neuromodulatory activity, whereas familiar or predictable patterns minimize prediction error and attenuate pupil-linked arousal.

Here, we bring these lines of research together to test how implicit auditory memory modulates pupil responses. Using a well-established paradigm, we induced memory for a set of regularly repeating auditory sequences that were unique to each participant. We then measured pupil dynamics while participants listened to previously encountered regular sequences and to novel regular sequences matched in structure.

We formulated two competing hypotheses. According to an **increased-response hypothesis**, familiar sequences should evoke larger pupil dilations, reflecting retrieval-related processes or enhanced salience, consistent with the pupil old/new effect. Alternatively, a **reduced-response hypothesis** predicts smaller pupil dilations for familiar sequences, reflecting more efficient processing and stronger predictability. Finally, by comparing active vs. incidental exposure listening conditions, we tested whether pupil-linked signatures of auditory memory emerge even in the absence of explicit attention to sound structure.

## Materials and methods

### Participants

**Group 1:** A total of 34 participants were recruited. Data from four participants were excluded from the analysis, leaving 30 participants in the final dataset (26 female, average age 25.2, range 19-32). Participants were excluded for being outliers in RT to STEP (see below) in the memory induction task (1 participant) or due to difficulty tracking eye data (3 participants; this occurred due to participant fatigue and/or excessive blinking). We aimed for N=30 in line with previous work (Huviyetli & Chait, 2026; Milne et al., 2021) that demonstrated robust, sustained pupil effects for such cohort sizes.

**Group 2:** A total of 54 different participants were recruited. Data from 14 participants were excluded from the analysis, leaving 40 participants in the final dataset (29 female, 26.27 average age, range 19-34). Participants were excluded due to poor performance in the gap detection task during memory induction (<50% performance; 7 subjects) or due to poor decoy task performance during pupillometry (1 subject), or due to technical issues during eye tracking (6 participants; this occurred due to participant fatigue and/or excessive blinking). We targeted a larger sample size in this group to account for the expected variability in performance arising from the decoy nature of the experimental manipulation.

All participants reported no hearing or neurological problems and provided written consent. Experimental procedures were approved by the UCL ethics committee.

### Group 1 Session structure and procedures

The experiment consisted of three stages:

**(1) Memory induction** using the ApMem (“Auditory Pattern Memory”) task as described in (Bianco et al., 2020). 40 min.
**(2) Pupillometry.** Participants listened to new (REGn) and previously encountered (REGr) REG patterns whilst performing a decoy gap detection task. 40 min.
**(3) Explicit familiarity.** A brief test to assess listener familiarity with REGr patterns. 5 min.

Participants were kept naïve to the purpose of the experiment and were not informed that patterns were reoccurring until the final explicit familiarity block.

#### Memory Induction

We used an implementation of the **ApMEM task** as described in (Bianco et al., 2020). Stimuli (Fig. 1A) were sequences of contiguous 50-ms tone-pips of different frequencies generated at a sampling rate of 44.1 kHz and gated on and off with 5-ms raised cosine ramps. Twenty frequencies (logarithmically spaced values between 222 and 2,000 Hz; 12% steps; loudness normalised based on ISO226) were arranged in sequences with a total duration of 6s. The specific order in which these frequencies were successively distributed defined different conditions that were otherwise identical in their spectral and timing profiles. **RND** (‘random’) sequences consisted of tone-pips arranged in random order. This was implemented by sampling uniformly from the pool with the constraint that adjacent tones were not of the same frequency. Each frequency was equiprobable across the sequence duration. The **RNDREG** (random-to-regular) sequences contained a transition between a random (RAN), and a regularly repeating pattern: Sequences with initially randomly ordered tones changed into regularly repeating cycles of 20 tones (an overall cycle duration of 1 s; new on each trial). The change occurred randomly between 2.5 and 3 s after sequence onset. **RND** and **RNDREGn** (RANREG novel) conditions were generated anew for each trial. Additionally, and unbeknownst to participants, 6 different REG patterns reoccurred identically within each block (**RNDREGr** condition, reoccurring). The RAN portion of RNDREGr trials was always novel. Each of the 6 regular patterns (REGr) reoccurred 3 times per block (every ∼ 2 minutes, i.e., 12 presentations overall). Reoccurrences were distributed within each block such that they occurred at the beginning (first third), middle, and end of each block. Two control conditions were also included: sequences of tones of a fixed frequency (CONT), and sequences with a step change in frequency partway through the trial (STEP). The STEP trials served as a lower-bound measure of individuals’ RT to simple acoustic changes.

**Figure 1:**
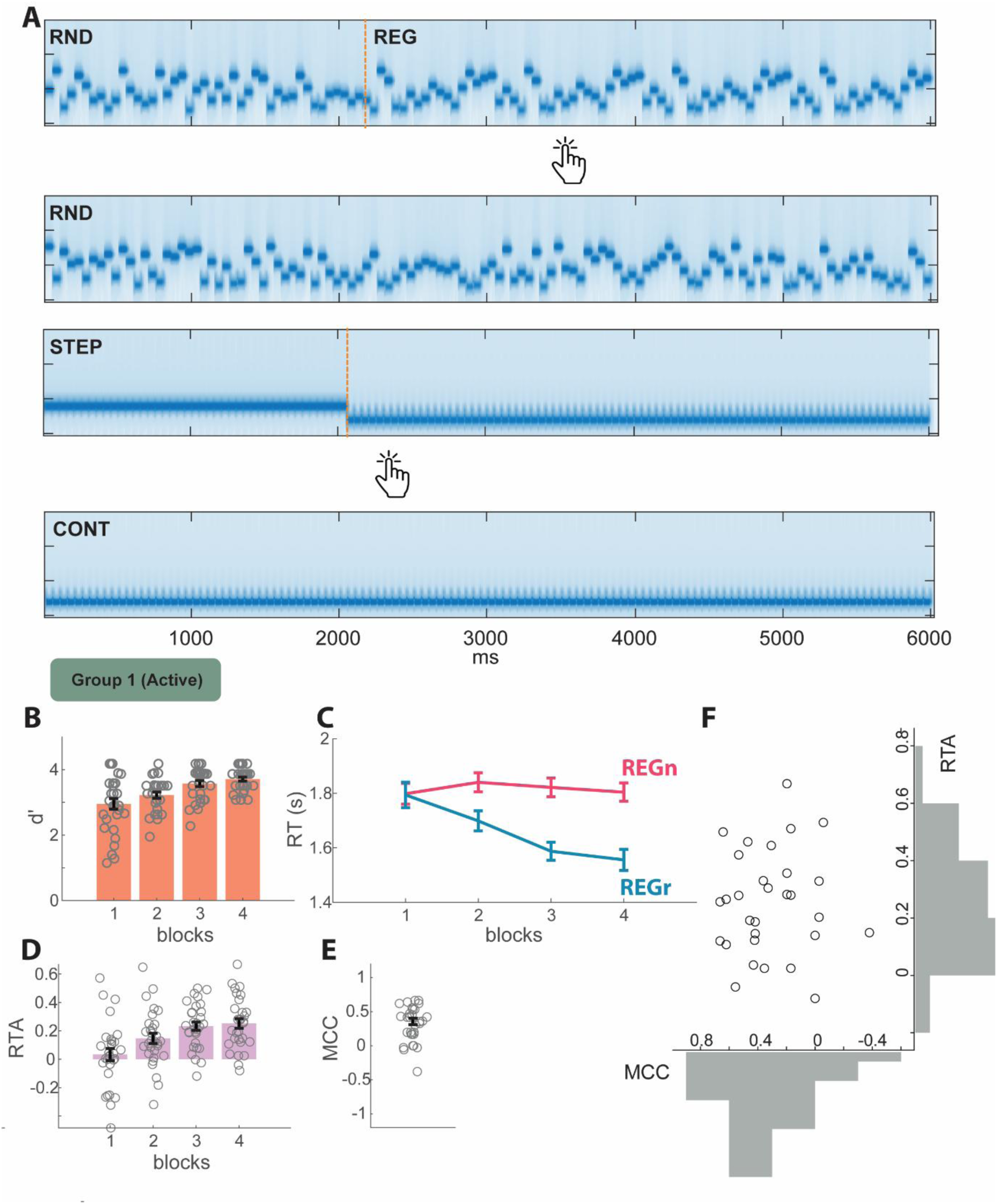
Group 1 behavioural results demonstrate the implicit formation of memory for REGr. **A** Spectra of example stimuli. Participants monitored unfolding tone sequences for the emergence of a regular pattern (in RND-REG stimuli) or a step change in frequency (in STEP stimuli) and were instructed to respond by button press as quickly as possible upon detection. **B** d’ sensitivity index in the main task (computed over REGn and REGr trials). Error bars indicate 1 STDE; individual participant data are indicated with grey circles. **C** (corrected) reaction time. **D** Reaction time advantage computed as the difference in reaction time between REGn and REGr. Error bars indicate 1 STDE; individual participant data indicated with grey circles. **E** MCC across participants (grey circles). **F** MCC vs. RTA for individual participants (circles).

Sounds were delivered with headphones (Sennheiser HD558). Participants were instructed to monitor for transitions from random to regular patterns (RNDREG) and frequency changes in STEP stimuli and press a keyboard button as soon as possible upon pattern detection. To acquaint participants with the task, a practice run was administered. The main task consisted of 4 blocks. Each block lasted about 8 minutes and contained 60 stimuli (18 RND, 18 RNDREGn, 18 RNDREGr, 3 STEP, 3 CONT), with an inter-trial-interval randomized between 0.5-0.8 s. Participants were instructed to respond as quickly and accurately as possible both to the transition from random to regular pattern and to the step frequency change. Feedback on accuracy and speed was provided at the end of each trial, as in our previous work (Bianco et al., 2020): a red cross for incorrect responses, and a tick after correct responses. The colour of the tick was green if responses were ‘fast’ (< 2200 ms from transition to REG or <500 ms from the step frequency change), and orange otherwise.

d’ (computed per block across RNDREGn and RNDREGr conditions) served as a general measure of sensitivity to regularity. Responses were marked as hits when they occurred after the pattern began to repeat (i.e., after the first cycle; effective transition). Responses that occurred earlier or during random trials were marked as false alarms. The core analysis focused on the RTs to the onset of regular patterns, as in (Barascud et al., 2016; Bianco et al., 2020). RT was defined as the time between the onset of the regular pattern (or the frequency step change) and the participant’s button press. For each participant and block, RTs that deviated more than ±2 standard deviations from the participant’s median RT were excluded. We then calculated the mean RT for both REGn and REGr trials within each block. For REGr trials, the mean was computed across the three intra-block presentations of the six distinct REGr patterns. Finally, to quantify memory formation, we computed the reaction time advantage (RTA) for each participant and block by calculating the difference in mean RTs between REGn and REGr trials.

#### Pupillometry

Only REGr and REGn sequences were presented. REGr were the recurring patterns from the memory induction task. REGn were newly generated for each trial. Since pupil responses tend to be slow, we used longer sequences (9 seconds) in line with previous work (Milne et al., 2021). The task consisted of 6 blocks (∼4 min each). Six REGr and 6 REGn were included in each block. Additionally, a total of 1 to 6 REGn stimuli that contained a silent gap between 1 and 8s post-onset were introduced. The number of gap-containing trials was varied across blocks to reduce predictability and prevent participants from forming expectations about a fixed number of such trials. The gaps were created by removing 3 tones (150ms) from each sequence. In each block, the number of trials ranged from 13 to 18 (6 REGr, 6 REGn, and 1 to 6 REGn with gaps). The order of stimuli was random with an inter-stimuli interval of ∼5 sec. Therefore, participants encountered each REGr pattern 6 times during the pupillometry experiment, and the presentations were spaced at least 10 min apart (block duration + break).

Participants sat with their head fixed on a chinrest in front of a monitor (24-inch BENQ XL2420T with a resolution of 1920x1080 pixels and a refresh rate of 60 Hz), in a dimly lit and acoustically shielded room (IAC triple-walled sound-attenuating booth). The distance between the chinrest and the screen was 62 cm. Sounds were delivered diotically to the participants’ ears with Sennheiser HD558 headphones (Sennheiser, Germany) via a Roland DUO-CAPTURE EX USB Audio Interface (Roland Ltd, UK), at a comfortable listening level (self-adjusted by each participant). Stimulus presentation and response recording were controlled with Psychtoolbox (Psychophysics Toolbox Version 3; Brainard, 1997) on MATLAB (The MathWorks, Inc.). Participants were instructed to fixate on a continuously presented fixation cross (black; on a grey background) and monitor the auditory stimuli for occasional silent gaps, which required a speeded button press. Feedback on accuracy was provided. A summary on accuracy, including the total number of false alarms, misses, and correct responses, was also given at the end of each block. A break of at least three minutes was given between blocks.

Pupil information was continuously recorded binocularly using an Eyelink 1000 Desktop Mount infrared eye-tracking camera (SR Research Ltd.) positioned under the monitor, focusing binocularly with a sampling rate of 1000 Hz. Before each experimental block, the standard five-point calibration procedure was carried out. Participants were told to blink naturally. They were also encouraged to rest their eyes briefly during inter-trial intervals. Prior to each trial, the eye-tracker automatically checked that the participants’ eyes were open and fixated appropriately; trials would not start unless this was confirmed.

#### Explicit familiarity

Explicit memory for REGr was examined with a surprise task at the end of the session. The 6 REGr patterns (only one instance per REGr) were intermixed with 36 REGn patterns. Participants were instructed to indicate which patterns sounded ‘familiar’. The classification was evaluated with the Matthews Correlation Coefficient (MCC) score (Boughorbel et al., 2017; Powers, 2007). Using the values of true positives (TP), true negatives (TN), false positives (FP), and false negatives (FN), the MCC score was computed using the formula 𝑀𝐶𝐶= 𝑇𝑃𝑋𝑇𝑁−𝐹𝑃𝑋𝐹𝑁√(𝑇𝑃+𝐹𝑃)(𝑇𝑃+𝐹𝑁)(𝑇𝑁+𝐹𝑃)(𝑇𝑁+𝐹𝑁). The MCC score ranges from 1 to -1, with 1 denoting perfect classification and -1 denoting perfect misclassification. An MCC score of zero indicates chance-level performance.

### Group 2 Session structure and procedures

The experiment consisted of three stages, with only the first stage differing from that of Group 1.

**(1) Incidental exposure with a decoy task (“Exposure”):** Participants listened to the same stimuli as Group 1; however, instead of performing an active pattern detection task, they engaged in a decoy gap detection task. This task required attention to the stimulus sequences but did not involve explicit detection of regular patterns. 40 min.
**(2) Pupillometry:** (identical to Group 1)
**(3) Explicit familiarity:** (identical to Group 1)

Participants were kept naïve to the purpose of the experiment and were not informed that patterns were reoccurring until the final explicit familiarity block.

#### Incidental exposure with a decoy task (“Exposure”)

The stimuli and presentation timing were exactly the same as those described for Group 1, except some of the trials contained a 50 ms gap replacing a single tone pip randomly chosen between 1 – 4.5 s post stimulus onset that participants were instructed to detect. Specifically, 5 of RNDREGn, 5 RND, and 1 STEP and 1 CONT trials contained a gap. None of the RNDREGr stimuli contained a gap to avoid any possible interference with memory formation. Therefore, as above, each block contained 60 stimuli, 12 of which contained a gap. Participants were instructed to fixate on a continuously presented fixation cross (black; on a grey background) and monitor the auditory stimuli for these occasional events which required a speeded button press. Feedback on accuracy was provided at the end of each trial. A summary on accuracy, including the total number of false alarms, misses, and correct responses, was also given at the end of each block. A break of at least three minutes was given between blocks.

For the decoy task, we aimed to maintain a high level of participant engagement - comparable to that of the regularity-detection task performed by Group 1, in which both speed and accuracy were emphasized, while keeping the sensory input identical across groups. To this end, we employed a challenging gap-detection task, in which participants were required to detect the removal of a single tone pip from the sequence. As expected, several participants were unable to perform this task (see above) and were excluded from further analysis. In contrast, the gap-detection task used during pupillometry was set at an easier level (removal of three tones), which most participants were able to perform with high accuracy.

### Analysis of pupil data

Trials containing a gap and trials with false alarms were excluded from the analysis. Most participants made infrequent false alarms and only one participant had more than one false alarm in a block. Blocks where the participant failed to respond to any gaps, were excluded from the analysis as we could not ensure that the participant was paying attention. This happened to data of only one participant (group1). Participants who failed to respond to 50% of the gaps were excluded from the analysis. This happened to one participant in group2.

#### Preprocessing

The left eye was analyzed whenever it was possible. Pupil data from each trial were epoched from 1s pre-onset to 10s post-onset. Intervals where the participant gazed away from fixation (visual angle > 2.56 degrees horizontal and 2.57 degrees vertical) or where full or partial eye closure was detected (e.g., during blinks) were automatically treated as missing data. Epochs with excessive missing data (>50%) were excluded from further analysis. For the rest, missing data were recovered with shape preserving piecewise cubic interpolation. Data were then smoothed with a Hanning window of 150ms. In group 1, a mean of 3 trials per subject (out of a total of 72) were removed (max =15; median=1.5; at most 10 trials in each condition). In group 2, a mean of 4.2 trials per subject (out of a total of 72) were removed (max =21; median=1.5; at most 13 trials in each condition). Following pre-processing, each participant’s pupil data (collapsed over REGn and REGr) were normalized (z-score) based on the baseline period, then averaged to create a single time series for each condition.

#### Statistical analysis

A non-parametric bootstrap approach was used to assess whether pupil dilation responses differed between the REGr and REGn conditions (Efron & Tibshirani, 1993; Milne et al., 2021). For each participant, a difference time series was computed by subtracting the REGr response from the REGn response. These individual difference time series were then subjected to bootstrap resampling (1,000 iterations with replacement). At each time point, a significant difference was identified if more than 95% of the bootstrap samples lay consistently above or below zero (i.e., p < 0.05). The analysis was performed across the full-time window shown in the plots, and all time points meeting the significance criterion are indicated.

To compare time-domain data between the two groups, we used a similar bootstrap approach in which, for each iteration, a number of subjects—equal to the size of the smaller group—was sampled with replacement from both groups, and the group means were then subtracted.

## Results

### Group 1: Participants developed an implicit memory of REGr patterns

Figure 1B presents the behavioural results from the memory induction stage. As expected, d’ values for regularity detection were high (Figure 1B), confirming that participants were able to reliably detect the emerging regular patterns—supporting the interpretability of the reaction time (RT) data. A repeated-measures ANOVA with block number as a factor revealed a main effect of block (F(1,29) = 38.4, p < 0.001, η² = 0.384), indicating a gradual increase in d’ across blocks. The RT data in Figure 1C show the emergence of a reaction time advantage (RTA), with responses to REGr patterns becoming increasingly faster than those to REGn patterns beginning in Block 2 and reaching a plateau in Block 3. This pattern is consistent with previous findings (Bianco et al., 2020, 2023). A repeated-measures ANOVA with factors of condition (REGr vs. REGn) and block revealed: a main effect of condition (F(1,29) = 47.8, p < 0.001, η² = 0.622), a main effect of block (F(3,87) = 6.5, p < 0.001, η² = 0.183), and a significant interaction between condition and block (F(3,87) = 13.4, p < 0.001, η² = 0.316). To explore the interaction, paired-sample t-tests were conducted for each block, comparing REGr and REGn RTs. These tests revealed: no significant difference in Block 1 (t(29) = 1.01, p = 0.92), and highly significant differences in all subsequent blocks (all p < 0.001).

This RT difference is summarized in the reaction time advantage (RTA) shown in Figure 1D. One-sample t-tests comparing RTA to zero confirmed: no significant RTA in Block 1 (t(29) = 0.77, p = 0.44), and significant RTAs in Blocks 2–4 (all p < 0.001; Cohen’s d > 0.74).

At the end of the session (after the pupillometry blocks), an explicit familiarity measure (expressed as an “MCC score”) was collected (Figure 1E). The mean MCC score across participants was 0.303, which was significantly above floor level (t(29) = 6.62, p < 0.001), suggesting some degree of explicit memory formation—albeit weak. This is consistent with previous findings (e.g., (Bianco et al., 2020)). Importantly, the MCC score did not correlate with the RTA in Block 4 (Figure 1F, Spearman’s ρ = –0.126, p = 0.500), suggesting that the RT advantage primarily reflects implicit memory.

Overall, these results confirm that participants implicitly acquired memory for the six reoccurring REGr patterns, in line with prior work (Bianco et al., 2020, 2023).

### Group1: REGr exhibits reduced pupil diameter relative to REGn

Figure 2 shows results from the pupillometry stage recorded while participants listened to previously experienced REGr patterns and novel, never-before-heard REGn patterns (Figure 2A). Performance on the incidental gap detection task (Figure 2C) was quantified as hit rate because false alarm rates were very low (Group 1: mean = 0.56 max=2 false alarms per subject; Group 2: no false alarms)E. Performance was high overall and did not differ between the two groups (Mann–Whitney U test *U* = 742, *p* = .0.323), suggesting participants were adequately engaged with the sounds.

**Figure 2:**
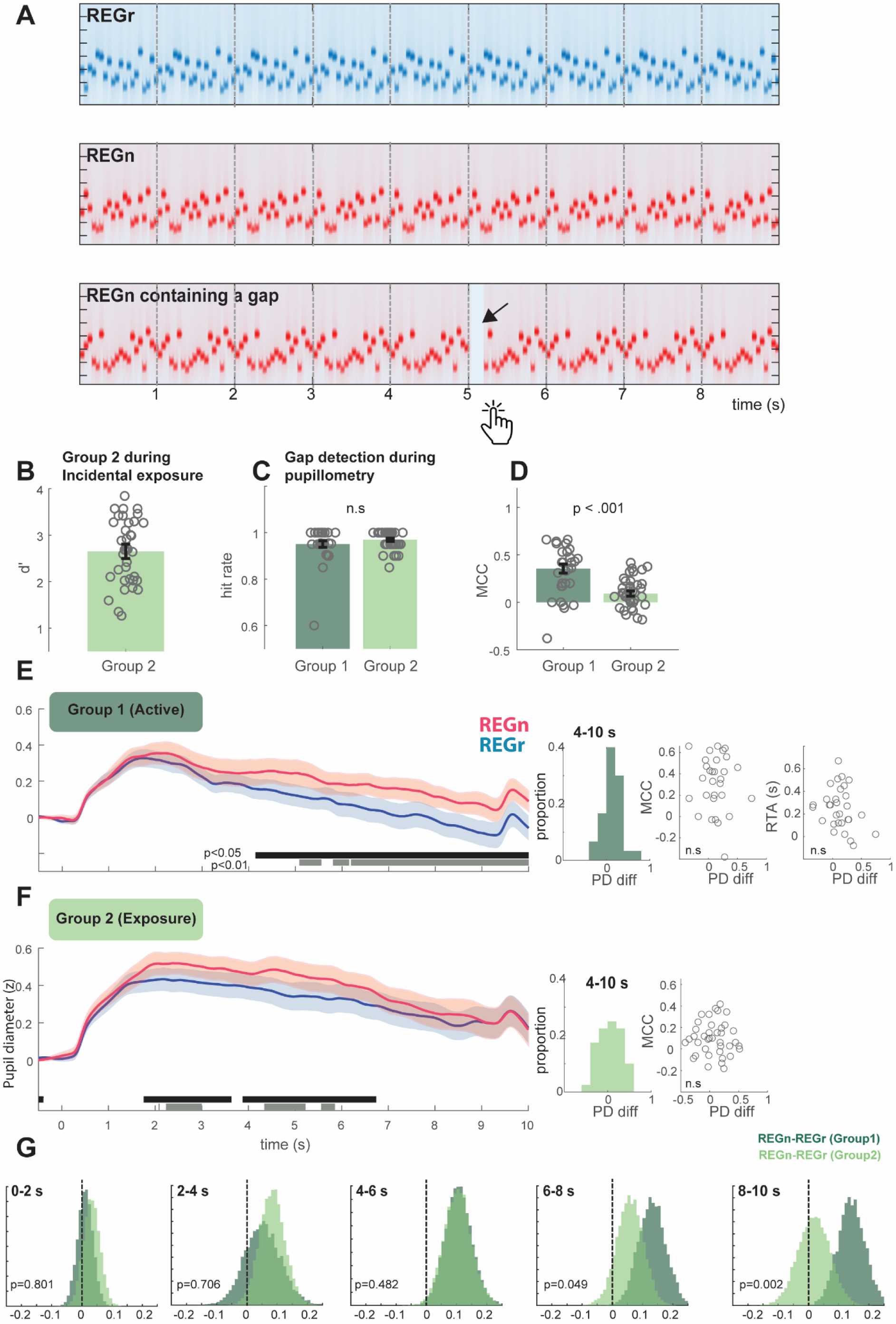
Pupillometry results reveal decreased pupil responses to REGr relative to REGn. **A** Spectra of example stimuli. Participants monitored 9 sec long REGn and REGr sequences for the presence of a silent gap (bottom; indicated with an arrow) and were instructed to respond by button press as quickly as possible upon detection. **B** Group 2: Gap detection performance during the incidental exposure phase (50 ms gaps). **C** Gap detection performance during pupillometry (150 ms gaps). Participants in both groups (Group 1 = dark green; Group 2 = light green) exhibited ceiling performance. **D** Familiarity performance (expressed as an MCC) score, obtained following the pupillometry blocks. Mean MCC scores were significantly different from 0 in both groups, and significantly higher in Group 1 relative to Group 2. **E** Pupillometry results from Group 1 showing time domain pupil size traces on the left. Grey and Black lines below the traces indicate significant differences; a distribution of differences (REGn-REGr between 4-10 s); and scatter plots PD difference against RTA and MCC (both non significant). **F** Pupillometry results from Group 2 showing time domain pupil size traces on the left. Grey and Black lines below the traces indicate significant differences; a distribution of differences (REGn-REGr between 4-10 s); and scatter plots of PD difference against MCC (non significant). **G** Resampled distribution of PD differences during different time intervals (indicated). p-values reflect the likelihood that the observed mean value for Group 2 came from the same population as Group 1.

We focus on the pupillometry results from group 1 first. Figure 2D presents the pupillometry data. The pupil responses exhibit a prototypical phasic dynamic, beginning approximately 300 ms post-onset, peaking initially around 500 ms, followed by a secondary peak around 2 seconds post-onset, and then gradually returning to baseline. A phasic response to the offset of the sequences is also observed.

Notably, during this post-onset period, REGr sequences elicited a faster reduction in pupil dilation, characterized by a smaller amplitude response that begins to emerge around 3 seconds post-onset, becomes statistically significant at 4 seconds, and persists until the end of the sequence. Figure 2D, right, also shows the distribution, across participants, of pupil dilation (PD) differences between REGn and REGr patterns, computed over the 4–10 second post-onset window. These PD differences did not correlate with either: the MCC score (Spearman’s ρ = 0.01, p = 0.958), or the RTA in Block 4 (Spearman’s ρ = –0.318, p = 0.087).

### Group 2: Incidental exposure also leads to pupil-linked memory for REGr (albeit weaker)

Throughout all stages of the experiment, Group 2 was exposed to the same auditory stimuli as Group 1. The key difference occurred during the initial memory induction phase: while Group 1 performed an active regularity detection task, Group 2 listened to the same sequences but engaged in a gap detection task instead. This task required sustained attention but did not involve explicit processing of regularities. Gaps consisted of the removal of a single 50 ms tone, making the task moderately challenging and attentionally demanding. The distribution of performance is shown in Figure 2B.

Subsequently, both groups proceeded to the pupillometry session, where they performed an easier version of the gap detection task involving 150 ms gaps. During the pupillometry blocks, each REGr sequence was presented six times—once per block, with at least 10 minutes between repetitions. After completing the pupillometry session, we assessed explicit familiarity using MCC scores.

The mean MCC score in Group 2 was 0.1 (Figure 2C), significantly above chance level (*t*(38) = 4.06, *p* < .001; Cohen’s *d* = 0.65), indicating that even participants in the incidental listening condition developed some explicit familiarity with the reoccurring REGr patterns. *(Note: Data from one participant in Group 2 are missing due to a technical error).* However, Group 2’s MCC scores were significantly lower than those of Group 1 (*t*(67) = 4.20, *p* < .001; Cohen’s *d* = 1.03; Figure 2C), suggesting that active engagement during the initial session, the only procedural difference between the groups, substantially enhanced explicit recognition of the reoccurring patterns.

Pupillometry results are shown in Figure 2E. A significant difference between the REGr and REGn conditions emerged approximately 2 seconds after trial onset and persisted until around 7 seconds post-onset—about 2 seconds prior to the offset of the regular sequence. Figure 2E, right, also shows the distribution, across participants, of pupil dilation (PD) differences between REGn and REGr patterns, computed over the 4-10 sec post onset window used for Group1. These PD differences did not correlate with either: the MCC score (Spearman rho= -0.51 p=0.76), or the performance in the memory induction task (Spearman rho= 0.16 p=0.4).

Overall, the pupil data in Group 2 indicate the presence of a pupil-linked implicit memory effect; however, this effect appears weaker in this group. To more closely examine differences between the two groups, we conducted a bootstrap resampling analysis (Figure 2F). For each participant in each group, we calculated the mean pupil difference between REGn and REGr within a defined time interval, then resampled these individual differences to generate a distribution for comparison (Figure 2G). The distribution for Group 1 (dark green) progressively shifts to the right of zero, indicating the build-up of a significant effect that stabilises around 4 sec post onset. Group 2 (light green) initially displays a similar pattern, but the distribution then shifts leftwards after 6 s post onset — where a group difference begins to emerge. Possible explanations for the early emergence and subsequent decline of the effect in this group are considered in the discussion below.

### MCC (explicit recognition) does not correlate with pupil effects

MCC scores (reflecting explicit recognition) did not correlate with pupil responses. To examine this further, participants were divided into high and low MCC groups using a median split within each group. As shown in Figure 3B-C, there was no significant difference in the pupil response (REGn minus REGr) between participants with higher versus lower MCC scores. This indicates that the observed pupil effects are not driven by explicit recognition of the REGr patterns during the pupillometry measurement, but likely reflect implicit memory processes.

**Figure 3:**
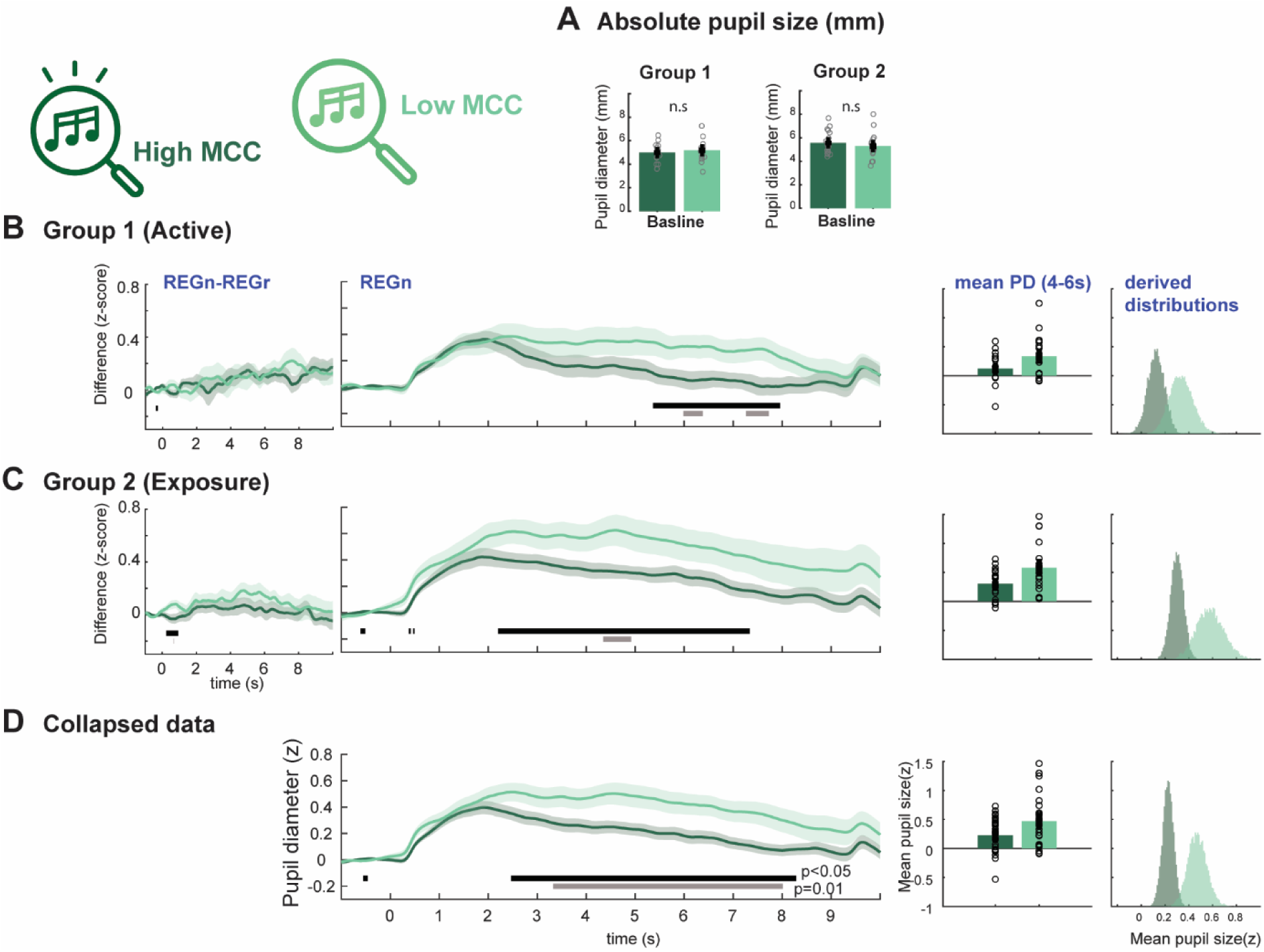
MCC (explicit familiarity) does not correlate with pupil-linked memory effect, but participants with high MCC show a reduced pupil diameter overall. **A** No difference in absolute pupil diameter between high/low MCC participants. **B** Group 1: Left: pupil-linked memory effect (REGn-REGr) is not different between high/Low MCC participants. Center: pupil response to REGn is different between high/low MCC participants. Right: values for individual participants and resampled distributions. **C** Group 2: Left: pupil-linked memory effect (REGn-REGr) is not different between high/Low MCC participants. Center: pupil response to REGn is different between high/low MCC participants. Right: values for individual participants and resampled distributions. **D** Collapsed data (Groups 1, 2) Center: pupil response to REGn is different between high/low MCC participants. Right: values for individual participants and resampled distributions. Grey and Black lines below the traces indicate significant differences.

### Participants with high MCC show reduced sequence-evoked pupil responses

An exploratory analysis was conducted to examine whether pupil responses differed between participants with high vs low MCC. Figure 3B-C-D compares responses to REGn between the high vs low MCC participants. Consistent across both Group 1 and Group 2, high-MCC participants exhibited a more pronounced reduction in pupil size over the course of the trial, compared to those with low MCC. This pattern was evident for both REGr and REGn conditions individually (see Supp Fig1). We chose to focus the main analysis on REGn, as it allows a more straightforward interpretation: these sequences consisted of novel, previously unencountered patterns and thus had equivalent status across groups. In Figure 3B-C-D, these effects are shown on the full time series traces, as well as per participant during 4-6 s. A univariate ANOVA with pupil diameter as the dependent factor, and MCC (high/low) and experiment group (group1, 2) as between subjects factor showed a main effect of MCC (F(1,65)=7.13; p=0.01; η² = 0.099) and a main effect of group (F(1,65)=5.25; p=0.025; η² = 0.075) with no interaction, consistent with overall pupil responses being somewhat larger in group 2.

Given that reduced pupil size has been associated with the detection and processing of regularities (Huviyetli & Chait, 2026; Milne et al., 2021), the present finding may reflect a relationship between the formation of robust memory representations, underlying subsequent familiarity judgments, and a broader mechanism supporting the efficient processing of structured stimuli.

## Discussion

We used pupillometry to probe the interaction between implicit auditory memory and neuromodulatory control, and to adjudicate between competing accounts of how familiarity shapes arousal regulation. Clarifying this relationship is important for both theoretical and practical reasons. Theoretically, it informs how the brain integrates past experience with ongoing sensory input to regulate arousal. Practically, it opens a path toward sensitive, task-free physiological markers of memory function, valuable in clinical and applied contexts where explicit behavioral measures may be unreliable or unavailable.

The pupil provides a valuable physiological index of cognitive and neural processes. Pupil diameter is jointly regulated by the sympathetic locus coeruleus–norepinephrine (LC–NE) system and parasympathetic pathways, serving as a sensitive indicator of stimulus-linked arousal. The neural circuits controlling pupillary dynamics are closely connected to central pathways implicated in learning and memory (Clewett et al., 2020; Grujic et al., 2024; Kafkas, 2024), suggesting bidirectional interactions that give rise to a complex interplay between pupil responses and cognitive states.

Most prior studies of pupil-linked memory have recorded pupil responses during active learning or retrieval, confounding memory-related arousal with the cognitive demands of task engagement. As a result, the specific contribution of memory to pupil dynamics has remained difficult to isolate. Here, we address this gap by demonstrating that implicit auditory memory alone, in the absence of explicit task demands, is sufficient to modulate pupil responses.

### Reduced pupil responses to implicitly remembered patterns

Consistent with previous reports (Bianco et al., 2020, 2023), participants exposed to sparsely reoccurring regular patterns (REGr; presented approximately every 3 minutes) developed an implicit memory, evidenced by reduced reaction times to REGr sequences relative to novel regular sequences (REGn). This reaction-time advantage emerged progressively across the first nine presentations of each REGr (three blocks) and then reached a plateau. Also consistent with prior work, this behavioural effect did not correlate with subsequent familiarity judgments, supporting its interpretation as an implicit memory trace rather than explicit recognition.

Critically, the REGr and REGn patterns, as well as the random (RND) sequences presented alongside, were composed of the same 20 tones and differed only in tone order. Successful recognition therefore required the encoding of sequential structure (Harrison et al., 2020) rather than simple feature-based information. Previous work has demonstrated that such sequence-specific memory is robust and long-lasting, persisting for at least six months following initial exposure (Bianco et al., 2023).

In the present study, following the memory-induction phase, participants completed a pupillometry experiment in which they listened to REGr and REGn sequences whilst performing a decoy task. Reported results are based on averages across 36 REGn and 36 REGr trials, with each of six REGr patterns presented six times, approximately 12 minutes apart (once per block). The large temporal gaps between repetitions of the same REGr patterns allowed us to attribute the observed implicit memory primarily to listeners’ experience during the induction session, rather than to memory traces formed during the pupillometry phase.

In previous work comparing REG and RND sequences (Huviyetli & Chait, 2026; Milne et al., 2021), the latency at which pupil-diameter traces diverged was substantially delayed relative to latencies derived from brain-imaging measures. It was therefore concluded that pupil dynamics are unlikely to reflect the process of regularity detection per se, but instead index downstream consequences for arousal, reflecting the organism’s experience of a predictable (“safe”) versus unpredictable (“high-alert”) environment. A prudent working hypothesis is that the same interpretation applies here. Although we do not have a direct ground-truth estimate of the detection time for REGr versus REGn (e.g. from M/EEG), behavioural reaction times provide a reasonable upper-bound benchmark, suggesting that neural differentiation should arise no later than 1–2 seconds post onset.

A particularly notable aspect of the present results is that both REGr and REGn sequences were *predictable*, especially after exposure to multiple cycles over 9sec (1-s cycles repeated nine times). M/EEG studies have shown that predictability representations for REGn sequences emerge within the first two cycles (Barascud et al., 2016; Bianco et al., 2026; Harrison et al., 2020; Hu et al., 2024; Magami et al., 2025; Southwell et al., 2017). Here, we used regularity cycles of 20 tones, where predictability develops more gradually and is represented more weakly than in shorter patterns. It is therefore plausible that predictability continues to be refined across repeated exposures across cycles within a trial. Nevertheless, REGr patterns elicited a stronger and more persistent memory trace than REGn, remaining evident even across these longer trials.

This is, to our knowledge, the first demonstration of pupil responses associated with implicit auditory memory. Interestingly, the pattern we observed contrasts with findings from the established pupil old/new recognition literature, in which remembered (“old”) items typically elicit larger pupil dilations, often interpreted as reflecting memory retrieval or attentional engagement. In contrast, the present results indicate that implicit memory representations lead to reduced pupil responses, consistent with more efficient processing and reduced neuromodulatory drive. Within a predictive coding framework, REGr sequences are associated with higher precision than REGn sequences, resulting in minimal prediction error. This reduced prediction error would in turn be expected to attenuate neuromodulatory engagement, consistent with the interpretation of smaller pupil responses as an index of lower uncertainty during auditory processing.

### Pupil-linked memory without explicit task demands

The question of how implicit memory formation is influenced by concurrent task engagement remains a central issue in auditory learning research (Andrillon et al., 2017; Bianco et al., 2020). The present findings demonstrate that pupil-linked signatures of implicit memory were evident even in Group 2, where participants were not explicitly instructed to detect regularities but were nonetheless actively engaged with the auditory input through a gap detection task. This suggests that implicit memory formation can occur automatically, without the necessity of overt attentional focus on the regular structure.

Although both groups exhibited clear memory effects, their temporal dynamics differed markedly (Figure 2). In Group 1, the memory-related reduction in pupil size was sustained to sequence offset. In Group 2, by contrast, the effect appeared transient—emerging initially but subsiding in the later portions of the trial. This dissociation suggests that active task engagement enhances the stability of implicit memory representations. In the absence of such explicit attentional focus, as in Group 2, the underlying memory trace may be weaker or less accessible, allowing the neural representation of novel regular patterns (REGn) to gradually approximate that of the reoccurring patterns (REGr).

From a broader theoretical perspective, this pattern aligns with accounts proposing that attention modulates the expression—but not necessarily the formation—of implicit memory traces (Batterink & Paller, 2019; Bianco et al., 2020; Turk-Browne et al., 2005). While exposure alone may suffice for encoding sequential regularities, sustained attentional engagement may be required to stabilize and maintain these representations over time, leading to more persistent physiological correlates such as reduced pupil-linked arousal.

### Dissociation from explicit familiarity

We also collected explicit familiarity judgments at the end of the experiment to assess whether participants could consciously differentiate the REGr patterns from novel sequences. There was substantial variability in MCC scores, and the mean familiarity performance differed between the two groups. Given that the pupillometry session was identical across groups, these differences most likely originated from the initial memory induction phase, indicating that explicit familiarity was stronger in the group that had been actively engaged with sequence structure.

Interestingly, however, in neither group did MCC scores correlate with the pupil-linked memory (Figure 2), suggesting that the two measures capture distinct aspects of memory. This dissociation implies that the pupil effect is not driven by explicit recognition of the REGr patterns or by any perceptual process involved in consciously distinguishing REGr from REGn sequences. Instead, the pupil modulation appears to reflect an implicit, automatic component of memory expression that operates independently of explicit awareness.

### Individual differences in pupil responsivity

An interesting incidental finding was that participants with higher MCC scores exhibited overall reduced pupil responses to REG patterns. Specifically, these individuals showed a more pronounced pupil constriction than participants with lower MCC scores. A trivial explanation for this effect could be that it reflects differences in general arousal or attentional engagement across participants. However, several aspects of the data suggest this is unlikely. First, the difference observed was evoked, emerging several seconds after sound onset rather than as a baseline shift. This temporal profile indicates that the effect reflects stimulus-driven processing rather than a general arousal state. Second, MCC status was not associated with performance differences in the decoy task during pupillometry (high vs low MMC in Group 1: *F*(1,28) = 1.40, *p* = .247; Group 2: *F*(1,37) = 0.52, *p* = .476), nor with gap-detection performance during the memory-induction phase (Group 2: *F*(1,37) = 0.71, *p* = .405). These findings argue against differences in task engagement as the source of the effect.

Instead, the observed relationship between pupil responses and MCC likely reflects individual variability in the ability to process structured auditory sequences. This interpretation echoes previous findings, discussed above, showing that regular, as compared with random, auditory patterns elicit smaller pupil responses (Huviyetli & Chait, 2026; Milne et al., 2021), suggesting that the magnitude and time course of pupil reduction index an individual’s sensitivity to regular structure. Taken together, these results indicate that MCC scores are linked to differences in the efficiency with which listeners extract and process sequential regularities.

The discovery of regularity is inherently memory-dependent (Harrison et al., 2020; Hu & Chait, 2026), and thus differences in working-memory capacity or sequence processing bandwidth are expected to modulate how efficiently individuals encode and utilize regular patterns. Indeed, the current study employed relatively complex 20-tone sequences, which are known to tax auditory memory systems (Barascud et al., 2016; Bianco et al., 2020). Prior work has shown that brain responses to such complex regularities lag behind ideal-observer estimates (Barascud et al., 2016; Southwell et al., 2017), suggesting limitations in the temporal integration window or representational bandwidth. It follows that Individuals with greater capacity to encode and exploit sequential structure may therefore exhibit both stronger pupil attenuation and higher familiarity performance. This raises the intriguing possibility that pupillometry could serve as an objective physiological marker of individual differences in structural learning capacity, beyond explicit recognition.

Importantly, however, this relationship appears to be distinct from implicit long term memory per se, as the pupil-linked memory effects reported earlier were independent of explicit familiarity performance. Together, these findings suggest that sensitivity to structure and implicit memory expression are related but dissociable components of auditory learning each reflecting different facets of the brain’s capacity to extract and exploit regularity in the acoustic environment.

Overall, the present findings demonstrate that pupil dynamics provide a robust physiological marker of auditory sequence tracking across multiple time scales. They reveal that implicit memory, even in naïve listeners, can modulate pupil-linked arousal in systematic and informative ways. More broadly, the results suggest that predictability in auditory sequences—whether acquired incidentally or established during active sequence tracking—interacts with the pupil-linked arousal system, reflecting adjustments in arousal and attention as the brain adapts to structured regularities in the acoustic environment.

## Acknowledgements

This work was supported by a BBSRC project grant to MC. The funders had no role in the study design, data collection and analysis, the decision to publish, or the preparation of the manuscript.

## Data availability

The data reported in this manuscript are available at DOI: <will be populated after acceptance>

---

**Supplementary Figure 1.**
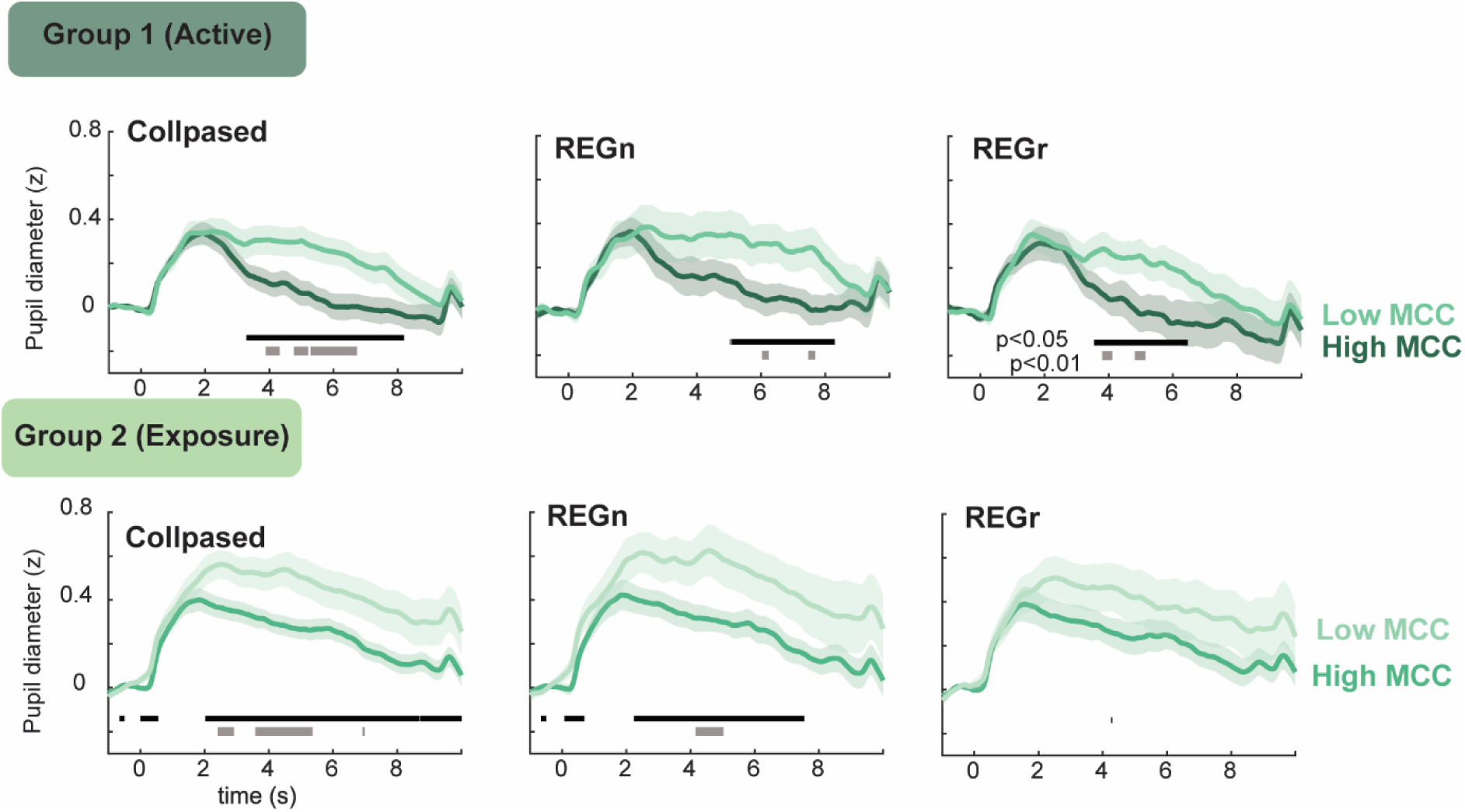
Participants with high MCC exhibit reduced sequence-evoked pupil responses. Shown are pupil response time courses for High- and Low-MCC participants, collapsed across conditions (REGn + REGr), as well as plotted separately for each condition. Horizontal lines indicate time periods with significant group differences. Note that minor differences in the timing of significant effects relative to Figure 3 reflect expected stochastic variability in the bootstrap analysis.

